# FIBOS: R and Python packages for analyzing protein packing and structure

**DOI:** 10.1101/2024.11.01.621530

**Authors:** Herson H. M. Soares, João P. R. Romanelli, Patrick J. Fleming, Carlos H. da Silveira

## Abstract

**Summary:** Advances in prediction of the 3D structures of most known proteins through machine learning methods have achieved unprecedented accuracies. However, although these computed structure models (CSMs) are overall remarkably good, they still challenge accuracy at the atomic level, especially in refinement of interatomic packing. The Occluded Surface (OS) algorithm is widely used for atomic packing analysis. But it lacks implementations in high-level languages like R and Python, limiting its accessibility. We introduce FIBOS, an R and Python package incorporating the OS methodology with enhancements. It embeds efficient Fortran code from the original OS. It also improves the original algorithm by using Fibonacci spirals to distribute surface dots, reducing anisotropy and ensuring a more even point distribution. As a case study using FIBOS, we compared packing densities between experimental protein structures and AlphaFold predictions. While average packing densities are similar, AlphaFold models exhibit comparatively greater variability and less homogeneity, potentially reflecting local inaccuracies. The FIBOS package provides an accessible tool for protein packing density analysis in R and Python, facilitating structural assessment in the era of extensive computational protein models.

**Availability and implementation:** The R and Python packages code are available at https://github.com/insilico-unifei/fibos-R and https://github.com/insilico-unifei/fibos-py.

## Introduction

Advances in protein structure prediction by machine learning methods have made estimating the 3D shape of virtually all known protein sequences a reality, with unprecedented accuracy (Jänes and Beltrao 2024). However, these advances introduce new challenges in low-level accuracy, especially the fine-tuning of the atomic positions (Bhattacharya and Cheng 2013, Binbay et al. 2023).

Molecule packing density calculations, in particular, play a crucial role in the structural analysis and assessment of protein models (Fleming and Richards 2000). Accurate packing density reflects the efficient organization of atoms, correlating directly with structural stability, functional integrity, and the realistic representation of interactions. Discrepancies in packing density can indicate potential inaccuracies, such as misaligned side chains, unrealistic voids, or incorrect folding, which can compromise the functional predictions of the model (Sonavane and Chakrabarti 2008). They are also important for algorithms that depend on plausible atomic coordinates for building reliable contact networks (da Silveira et al. 2009).

To date, one of the best-known algorithms for atomic packing analysis is Occluded Surface (OS) (Pattabiraman et al. 1995). This method distributes dots (representing patches of area) across the atom surfaces. Each dot has a normal that extends until it reaches either a van der Waals surface of a neighboring atom (the dot is considered occluded) or covers a distance greater than the diameter of a water molecule (the dot is considered non-occluded and disregarded). As a consequence, buried and well packed atoms, such as those in the protein core or in hot spot regions at chain-chain interfaces, tend to accumulate more dots on their surface and shorter normal lengths (Figure 1). On the other hand, those that are less packed, like near cavities or unstructured regions, may have fewer dots and longer normals. Thus, with the summed areas of dots and the lengths of normals, it is possible to compose robust metrics capable of inferring the average packing density of atoms, residues, proteins, as well as any other group of biomolecules.

**Figure 1.**
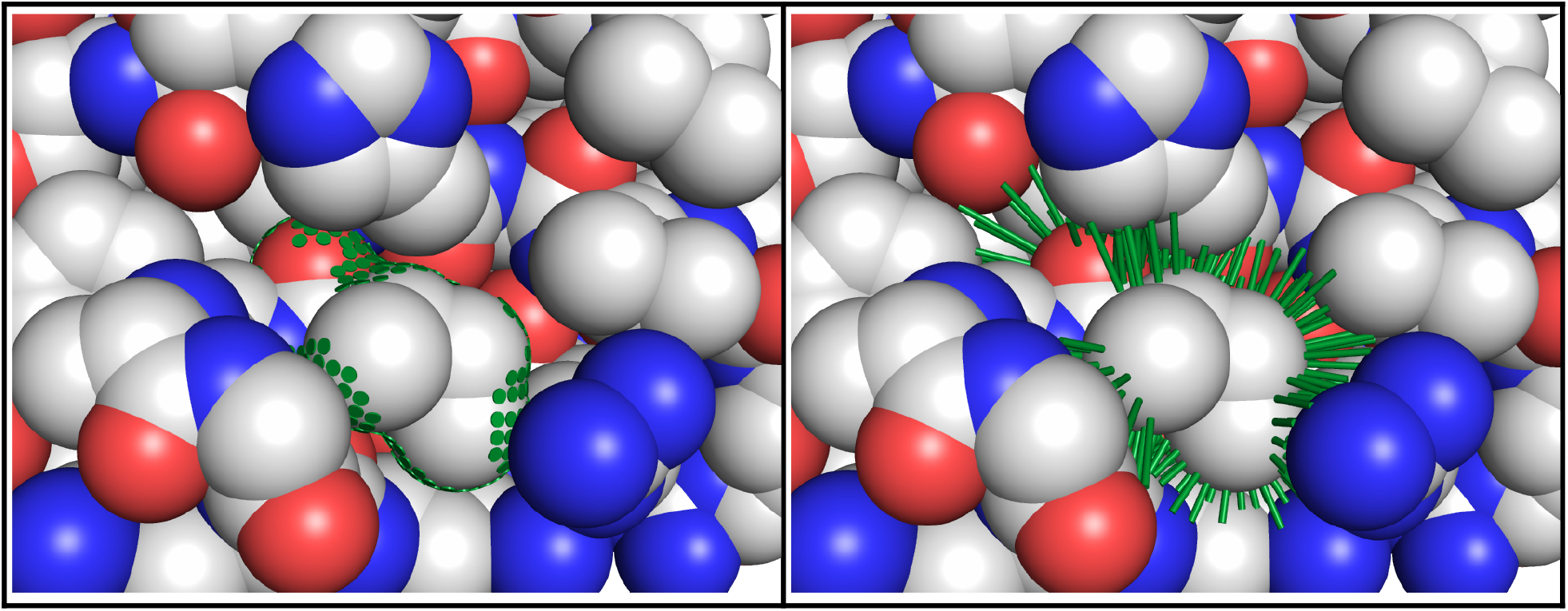
Dots (left) and normals (right) calculated with FIBOS for ILE 44 of ubiquitin 1UBQ. Partially exposed on the surface, this residue is a central part of a hydrophobic patch, critical for the binding of ubiquitin to other proteins. Pymol (DeLano 2015) was used to create the images using scripts included in the FIBOS distribution.

However, there are still no OS implementations in higher-level languages, such as R and Python. Both are fast and efficient coding languages with a wide user base across common operating systems and are especially suited to handling large volumes of data and analysis. Given that hundreds of millions of protein structure predictions are being generated by machine learning systems (Callaway 2022), having atomic packing density checking algorithms like OS in R and Python is a useful addition to available analytical tools.

Here, we present FIBOS, a package for biomolecule packing estimation in R and Python. It has embedded the same efficient Fortran implementations from the original OS, but extended with some improvements. There are built-in functions that return useful data in tables, like dots and normals by atom or residue. Also, a function that implements the occluded surface packing density metric (OSP) averaged by residue, as described in (Fleming and Richards 2000). Another important advancement concerns the algorithm for generating dots onto the surface. The original OS covered the surface radially with one of the axes as a reference, introducing not only axial anisotropies but also inhomogeneities in the dot areas when comparing poles and equator. In this version, Fibonacci spirals were used to allocate the dots, which is known to produce lower axial anisotropy as well as more evenly spaced points on a sphere (Swinbank and Purser 1999). Some comparative analysis between OS and FIBOS can be seen in Figures S1a, S1b, S2 and S3.

## Material and Methods

The user can choose between classical OS or new FIBOS (default) methodologies. The FIBOS package is multi-platform and runs on Linux, Windows and Mac. With appropriate code manipulations, FIBOS functions can perform parallelism across cores or threads, for example. Instructions and specifics for each operating system for R or Python can be found on the respective GitHub repositories. Fortran original code was retrieved from https://pages.jh.edu/pfleming/sw/os.

The dataset containing 97 PDB IDs of unique chains of the original OS publication (Fleming and Richards 2000) was used for package validation in R and Python compared to Fortran. The dataset can be seen in Table S5.

The `occluded_surface` and `osp` functions were used to calculate the OS at the atomic and residue levels per PDB ID, respectively. By default, these functions ‘clean’ the input pdb file by eliminating alternative conformations, heteroatoms and hydrogens. Atomic radii used are listed in Table S6.

## Results

To validate the libraries in R and Python, the average OSP of the full dataset mentioned above was calculated and the results compared with pure Fortran. Table S5 shows the OSP values in these 3 programming languages and Figure S4 presents the linear regression between them, revealing a high correlation above 0.99. This attests to the reliability of the values calculated in R and Python.

To evaluate the potential use of the new libraries in R and Python, a case study was set up, aiming to compare the packing density between experimentally determined structures and the same structures predicted by AlphaFold (AF). The R code for this study and supplementary materials can be found in GitHub repository (see Data Availability section).

The RCSB RESTful API (1.47.5) was used to download metadata from PDB IDs of Table S5, especially the Uniprot IDs and primary sequences (Rose et al. 2021). The Uniprot IDs served for composing the URLs needed to download the predictive models from the AF website (Varadi 2022). In this way, each computed structure model (CSM) of AF was associated with the respective experimental PDB. The primary sequence was used to select a subset of PDBs and respective CSM-AFs, so that the disparity in the number of residues and sequence alignment was not greater than 5 residues, which resulted in 44 structures (Table S7). Of these, 43 had the global predicted Local Distance Difference Test (pLDDT) greater than 90.00 (considered very high), with the median being 97.24. The lowest value was 83.02 (considered confident).

As can be seen in Figure 2, the average OSP values are similar when considering experimental PDB and predicted AF, given the similar profile of central tendencies and density distributions, with high p-values (Wilcoxon test = 0.087; Kolmogorov-Smirnov test = 0.99) and small Cohen’s d effect size (0.071). The same is not true for the standard deviations, revealing dissimilar points of central tendency and density distributions, with low p-values (Wilcoxon and Kolmogorov-Smirnov test < 0.001) and large Cohen’s d effect size (0.94). This indicates that the predicted AF models, when compared with the experimental ones, may exhibit greater variability and less homogeneity, potentially reflecting local inaccuracies (see Table S8).

**Figure 2.**
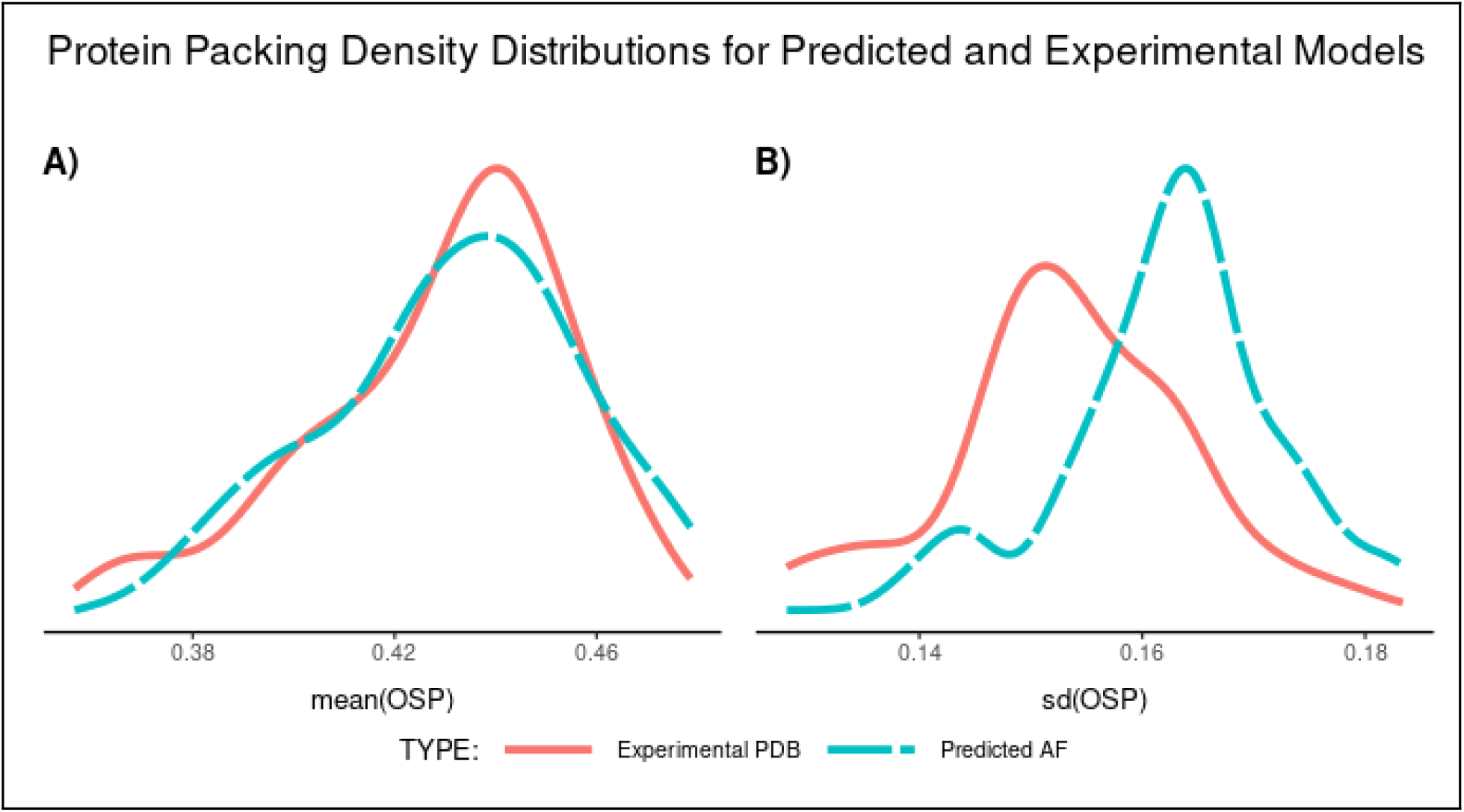
Protein packing density distributions involving 44 protein structures of Table S7, with experimental PDB (solid red lines) and predicted AF (dashed blue lines). A) The density distributions of whole protein average residue OSP values. B) The density distributions of the standard deviation of residue OSP values. See Table S8 for the statistical values.

## Conclusion

We present here the FIBOS package for R and Python, capable of calculating the atomic packing density in proteins. The ease and speed of coding that such high level languages provide is fundamental for the agility of analyses by researchers in structural biology, especially at this time of profuse data coming from computed structure models (CSM), such as AlphaFold (Varadi 2022), ESMfold (Lin et al. 2023) and RoseTTAFold (Krishna et al. 2024).

We show a brief case study to illustrate how FIBOS can be used to analyze differences in atomic packing density between predicted and experimental protein structure models. This study suggests that there are differences, with the former presenting greater variability in packing densities than the latter. Further investigation will be needed to assess this bias in detail.

## Data availability

Supplementary data are available at Bioinformatics online. Data used for package validation and case study in: https://github.com/insilico-unifei/fibos-R-case-study-supp.

## Conflict of Interest

None declared.

## Funding

This work has been supported by FAPEMIG.

## Notes

### Competing Interest Statement

The authors have declared no competing interest.

