## SUPPLEMENTARY MATERIALS for "FIBOS: R and Python packages for analyzing protein packing and structure"

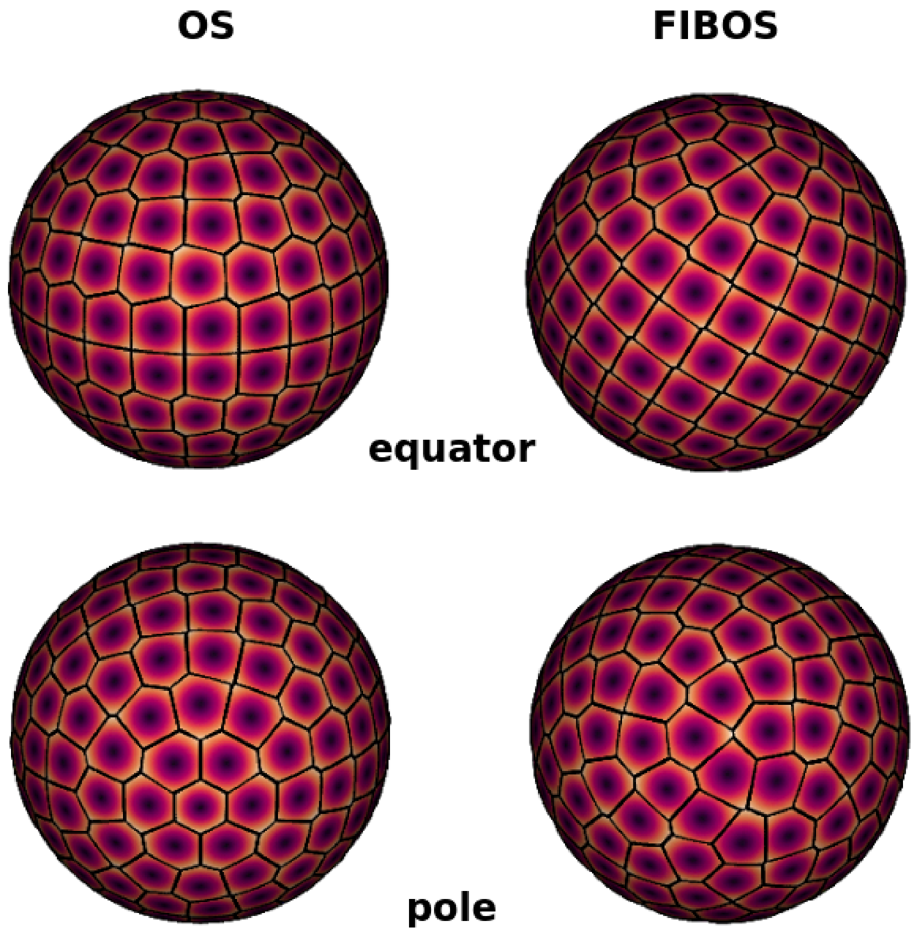


**Figure S1a.** Voronoi tiling on a sphere, comparing dot distributions for OS (left) and FIBOS (right). Above, it shows the equators; below, the poles. The spheres have a radius of 1.90 Å (equivalent to the van der Waals radius of carbon) and 212 dots were distributed, the default number used in the original OS.

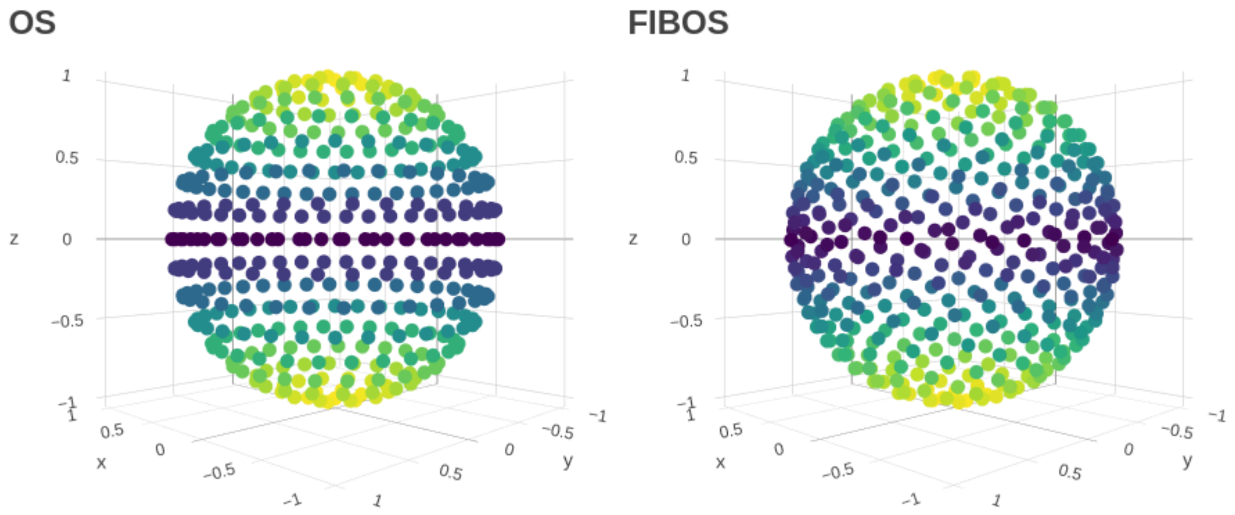


**Figure S1b.** Another way to see the Voronoi tiling on a sphere, for OS (left) and FIBOS (right). It is possible to see the anisotropy and bias in the distribution of points, more prominent in OS than FIBOS.

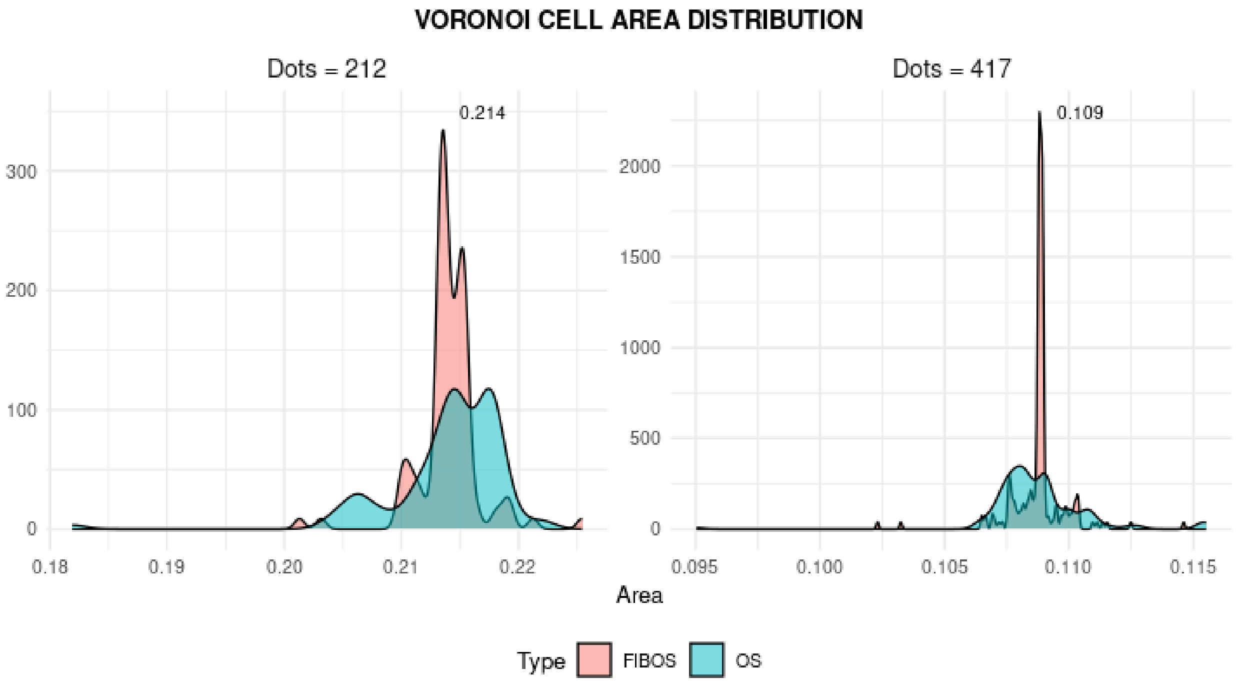


**Figure S2.** Voronoi cell area distributions, comparing FIBOS and OS with 212 and 417 dots on a spherical surface of radius 1.90 Å (equivalent to the van der Waals radius of carbon). The numbers near to the modals depict the mean of cell areas. FIBOS presents a profile of cell areas more concentrated around the mean value, indicating more evenly spaced points on a sphere.

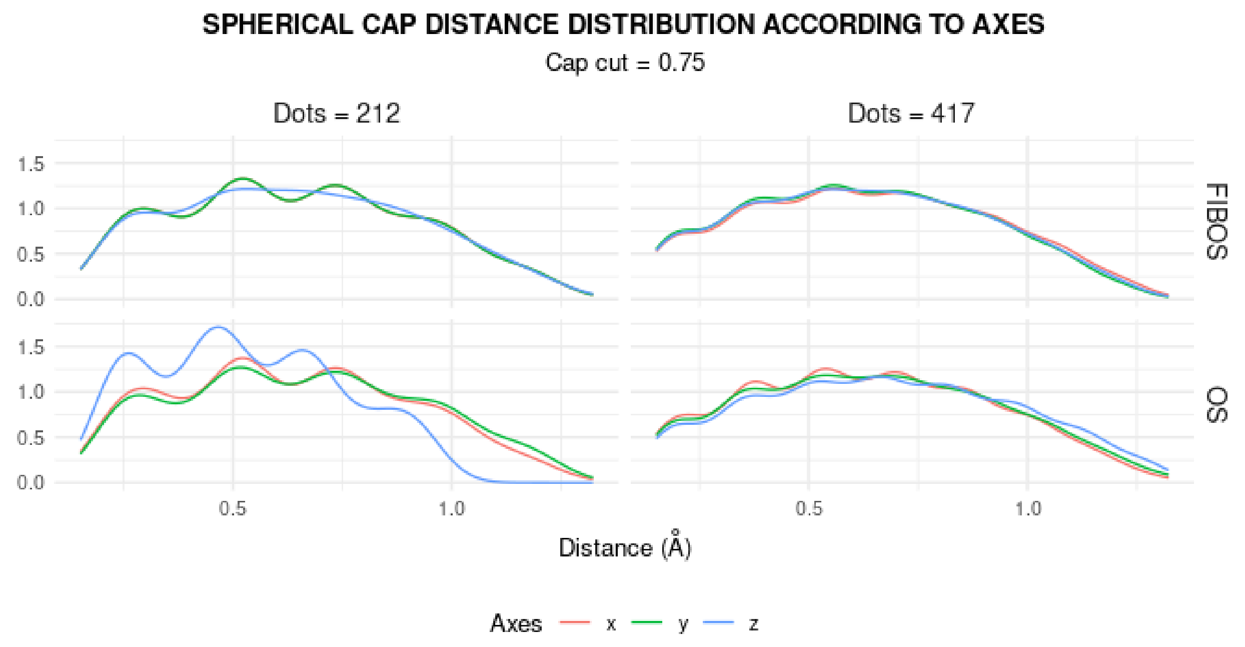


**Figure S3.** Distance distribution of all dots from each other, but limited to spherical caps at 0.75 of the radius, on the x, y and z axes. There is a bias in the distance distribution considering the spherical cap along the z axis, but it is much smaller in FIBOS than OS. So FIBOS presents lower axial anisotropy than OS, mainly for moderate amounts of dots.

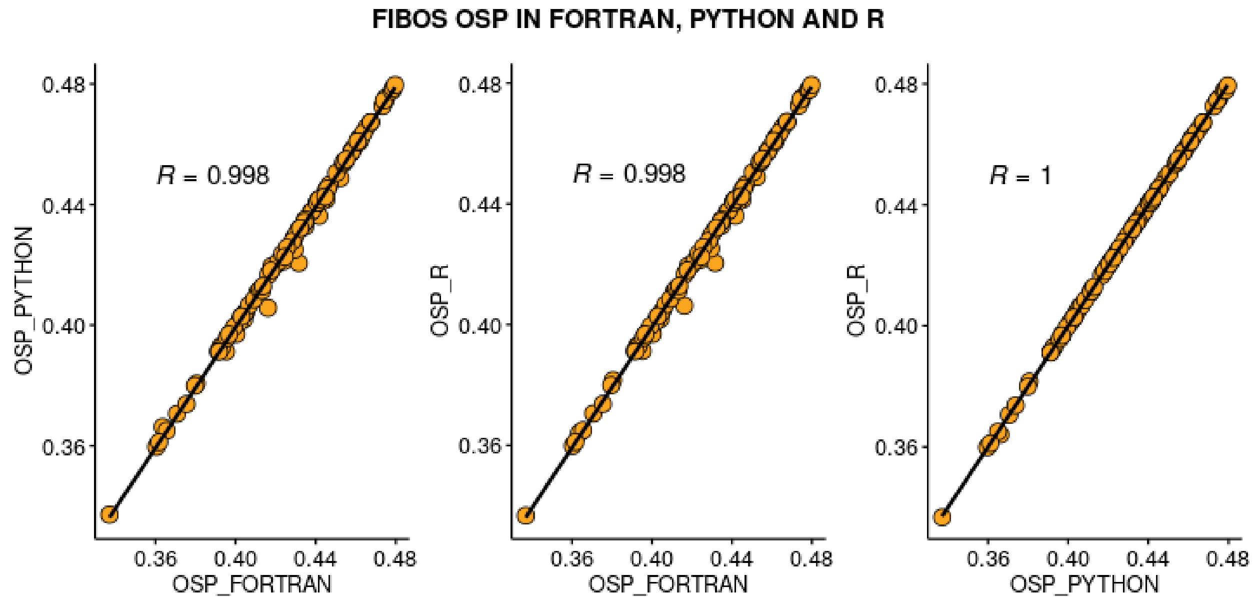


**Figure S4.** Linear regression indicating high correlations between OSP calculations between Fortran, R and Python, for dataset from Table S5.

**TABLE S5:** Protein dataset used in testing and validation.

SUPPLEMENTARY:dataset:FIBOS:2024

**TABLE S6:** Atomic radii adapted from https://pages.jh.edu/pfleming/sw/os.

SUPPLEMENTARY:radii:FIBOS:2024

**TABLE S7:** Subset of Table S5 used in study case.

SUPPLEMENTARY:study-case:FIBOS:2024

**TABLE S8:** average and standard deviation of residue OSP values of each entry of Table S7.

SUPPLEMENTARY:study-case-stat:FIBOS:2024
